# Defining Operational UV-C Dose Requirements for Autonomous Disinfection of Clinically Relevant Pathogens Across Healthcare and High-Touch Surfaces

**DOI:** 10.64898/2026.08.24.746724

**Authors:** Isaac Kong Fan Wu, Neel Ravi Vajaria, Luz Victoria Stam Viruega, Evelyn Wisebourt, Paul Fernanado Solis-Reyes, Kanghan Ryu, Elijah Ryan Ilasin, Amy Yang Shi, Nicole Jasmine Friesen, Khandaker Atkia Fariha, Stephen D. Barr

**Affiliations:** Department of Microbiology and Immunology, Schulich School of Medicine and Dentistry, Western University, 1151 Richmond St., London, ON N6A 5C1, Canada

**Keywords:** Ultraviolet-C (UV-C), environmental disinfection, healthcare-associated pathogens, UV-C dose-response, autonomous disinfection, high-touch surfaces

## Abstract

**Background:** Autonomous ultraviolet-C (UV-C) disinfection systems are increasingly used to supplement manual environmental cleaning, yet evidence-based guidance defining pathogen-specific UV-C dose requirements across representative surfaces remains limited.

**Aim:** To characterize operational UV-C dose requirements for clinically relevant pathogens across diverse high-touch and healthcare surfaces and determine how experimentally derived microbial inactivation can inform operational exposure parameters.

**Methods:** SARS-CoV-2, adenovirus, *Pseudomonas aeruginosa*, *Staphylococcus aureus*, *Klebsiella pneumoniae*, *Enterococcus faecalis*, *Candida auris*, and *Clostridioides difficile* spores were exposed to defined UV-C doses on representative high-touch materials or stainless steel under standardized conditions, including a 10% fetal bovine serum organic soil challenge. Microbial inactivation was quantified by viable recovery. Dose-response analysis and operational modelling were used where supported by the experimental data.

**Findings:** UV-C exposure significantly reduced viable recovery of all pathogens, with substantial differences in the exposure conditions associated with microbial inactivation. SARS-CoV-2 exhibited substantial inactivation at doses as low as 2.6 mJ/cm^2^, whereas the highest evaluated doses were 1,800 mJ/cm^2^ for *C. difficile* spores and 3600 mJ/cm^2^ for *C. auris*. For *C. auris*, multi-dose data estimated that approximately 1,410 mJ/cm^2^ was associated with a 2-log_10_ reference reduction, enabling distance-dependent exposure-time predictions.

**Conclusion:** Experimentally quantified UV-C exposures produced substantial microbial inactivation across diverse pathogen classes and surfaces. Integrating delivered dose with microbial reduction provides a quantitative framework for translating laboratory efficacy into operational parameters for autonomous UV-C disinfection.

## INTRODUCTION

High-touch environmental surfaces are recognized reservoirs for infectious microorganisms and play an important role in the transmission of healthcare-associated pathogens. Contaminated surfaces contribute to the persistence and spread of viruses, bacteria, fungi, and bacterial spores within healthcare facilities and other high-occupancy environments, increasing the risk of healthcare-associated infections and outbreak propagation [1–4]. Effective environmental decontamination is therefore an essential component of infection prevention and control strategies.

UV-C irradiation (200–280 nm) is a well-established antimicrobial technology that inactivates microorganisms primarily through photochemical damage to nucleic acids, including the formation of pyrimidine dimers that inhibit replication [5]. Owing to its broad-spectrum antimicrobial activity, UV-C has demonstrated efficacy against viruses, bacteria, fungi, multidrug-resistant organisms, and bacterial spores and is increasingly used to supplement manual cleaning and chemical disinfection in healthcare environments [6–9].

Although numerous studies have demonstrated the efficacy of UV-C disinfection, relatively few have established quantitative operational UV-C dose requirements across diverse clinically relevant pathogens and representative environmental surfaces. Most studies have focused on individual pathogen classes or standardized laboratory substrates such as stainless steel, glass, or plastic, which provide important mechanistic insights but do not fully reflect the diversity of high-touch materials encountered in healthcare and other built environments. Surface composition, porosity, reflectivity, and topology can influence UV-C dose delivery and microbial inactivation, yet comparatively few studies have systematically evaluated representative polymers, textiles, composite materials, and healthcare stainless-steel surfaces within a single experimental framework. Furthermore, published studies frequently evaluate individual pathogens or specific pathogen classes independently, making direct comparison of operational UV-C requirements across viruses, bacteria, fungi, and bacterial spores difficult. Consequently, there remains limited evidence integrating pathogen diversity, representative environmental materials, and delivered UV-C dose into a practical framework that can guide deployment of autonomous UV-C disinfection systems.

Using controlled stationary exposure conditions, we established pathogen-specific operational UV-C dose parameters for the autonomous RAY UV-C disinfection system against a diverse panel of clinically relevant pathogens, including enveloped and non-enveloped viruses, Gram-negative and Gram-positive bacteria, bacterial spores, and fungal pathogens. Testing was performed on a diverse range of representative high-touch materials together with healthcare-relevant stainless-steel surfaces to better represent the diversity and complexity of surfaces encountered in healthcare and other high-touch environments. By integrating dose-response analyses, where available, with experimentally validated operational exposure conditions, we established a quantitative framework linking delivered UV-C dose, exposure conditions, surface type, and microbial inactivation. This framework provides evidence-based operational guidance for deployment of autonomous UV-C disinfection systems in healthcare and other high-touch environments.

## METHODS

### Materials and Experimental Design

RAY is an artificial intelligence (AI)-driven fully autonomous UV-C disinfection robot designed for healthcare and other high-touch environments (Figure 1). The system integrates a compact mobile platform with intelligent navigation, environmental sensing, and optimized path planning to deliver consistent UV-C disinfection. Onboard algorithms continuously calculate UV-C dose (emitted at a wavelength of 254 nm) requirements to optimize positioning, exposure duration, and movement, while integrated sensors enable autonomous mapping, obstacle avoidance, and route execution. The platform also supports remote monitoring, analytics, and reporting. Mechanically, RAY incorporates a central UV-C tower designed to provide 360° environmental exposure within a compact footprint. For the present study, RAY was evaluated under controlled stationary exposure conditions at defined distances to enable reproducible quantification of UV-C dose delivery and pathogen-specific inactivation thresholds across multiple materials and organism classes. The UV-C bulbs were given a two-minute warm-up prior to testing. This approach minimized variability associated with autonomous movement, environmental geometry, and dynamic positioning, thereby facilitating standardized comparison of UV-C efficacy under well-defined conditions.

**Figure 1.**
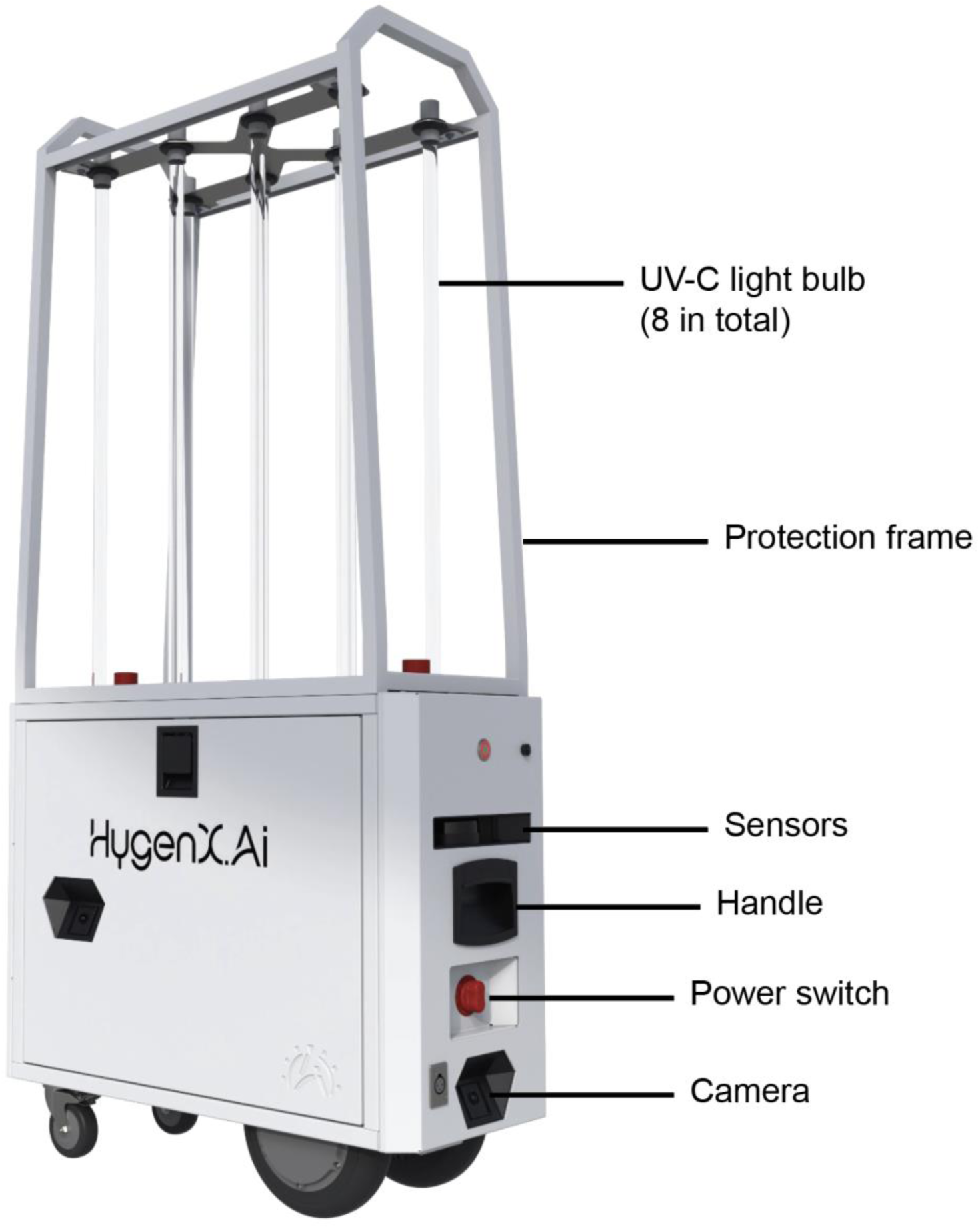
The RAY autonomous UV-C disinfection system. Photograph of the RAY AI-driven fully autonomous UV-C disinfection robot used in this study. The system combines intelligent navigation, onboard environmental sensing, and optimized path planning to deliver rapid, consistent, and repeatable UV-C room disinfection. RAY continuously performs real-time UV-C dose calculations to optimize positioning, exposure time, and movement, and features a central UV-C tower designed for 360° environmental exposure within a compact footprint.

A panel of materials representative of surfaces commonly encountered in healthcare and other high-touch environments was selected for UV-C disinfection testing. These included rigid thermoplastic polymers (polystyrene, Kydex, and Boltaron), cloth seating material, synthetic leather, and stainless steel, collectively representing surfaces commonly used in medical equipment, furnishings, seating, and other frequently touched environmental components. Material selection was guided by environmental relevance rather than exhaustive evaluation of all pathogen–surface combinations. Stainless steel was included as a standardized healthcare-associated surface because of its widespread use in clinical settings and established role in pathogen transmission. The material panel encompassed both smooth, non-porous surfaces (polystyrene, Kydex, and stainless steel) and more structurally complex surfaces (cloth, synthetic leather, and Boltaron) to evaluate UV-C performance across a range of surface characteristics that may influence microbial recovery and effective UV-C dose delivery through differences in surface texture, topography, and microshadowing.

### Pathogens

A panel of evolutionarily and clinically distinct pathogens was selected to evaluate the broad-spectrum antimicrobial efficacy of UV-C irradiation, encompassing multiple organism classes with known differences in structural complexity and UV susceptibility. Pathogens were tested under conditions of a 10% final soil load using fetal bovine serum (FBS). Initial inoculum concentrations were optimized for each organism and surface combination to produce recoverable untreated control counts within the quantifiable range of each assay (Supplementary Table S1), thereby enabling accurate determination of log_10_ reductions following UV-C exposure. To ensure active growth, all vegetative cultures were passaged daily for three consecutive days, with the final culture incubated for 48 h before experimentation.

SARS–CoV–2 (enveloped virus) isolates were acquired from BEI Resources, NIAID, NIH: SARS–Related Coronavirus 2, Isolate hCoV–19/USA–WA1/2020, NR–52281 (original isolate); SARS–Related Coronavirus 2, Isolate hCoV–19/South Africa/KRISP–EC–K005321/2020, NR– 54008, contributed by Alex Sigal and Tulio de Oliveira (beta variant); SARS–Related Coronavirus 2, Isolate hCoV–19/Japan/TY7–503/2021 (Brazil P.1, gamma variant), NR–54982, contributed by National Institute of Infectious Diseases. All viral work was performed under Containment Level 3 (CL3) conditions at the ImPaKT facility (Western University, Canada). Viral stocks were propagated in Vero E6 cells, a highly permissive African green monkey kidney cell line, maintained in Dulbecco’s Modified Eagle Medium (DMEM) supplemented with 10% FBS, penicillin and streptomycin. Cells were infected at approximately 50–60% confluency and incubated at 37°C with 5% CO_2_ for 72 hours. Supernatants containing infectious virus were harvested, clarified by centrifugation (500 × g, 10 min), aliquoted, and stored at −80°C. Human adenovirus type 5 (Adenoid 75; ATCC VR-5) (non-enveloped virus) was propagated in HEK293T cells under similar culture conditions. Following infection, cells and supernatants were collected after 72 hours and subjected to repeated freeze–thaw cycles to release intracellular virions. Lysates were clarified by centrifugation and stored at −80°C. Viral stocks were quantified prior to experimentation, with titers ranging from approximately 10^5.8^ to 10^7.5^ TCID50/mL.

*Staphylococcus aureus* (*S. aureus*) (ATCC 11632) was cultured in ATCC Medium 3 (nutrient agar/broth), and *Pseudomonas aeruginosa* (*P. aeruginosa*) (ATCC 15442) was cultured in ATCC Medium 18 (tryptic soy agar/broth). *Klebsiella pneumoniae* (*K. pneumoniae*) (BEI Resources NR-41899; strain BWH22), a carbapenem-resistant sequence type 258 (ST258) clinical isolate carrying blaKPC-2 and blaTEM-1, and *Enterococcus faecalis* (*E. faecalis*) (BEI Resources NR-31972; strain SF28073), a multidrug-resistant clinical isolate resistant to erythromycin, gentamicin, and vancomycin, were selected as representative healthcare-associated bacterial pathogens. *Clostridioides difficile* (*C. difficile*) (BEI Resources NR-13427), a multidrug-resistant clinical isolate resistant to fluoroquinolones, cephalosporins, and lincosamides, was included as a clinically important spore-forming healthcare-associated pathogen capable of persistent environmental contamination and transmission from contaminated surfaces. Vegetative cells and spores were prepared under anaerobic conditions as previously described [10]. Briefly, spores were germinated on pre-reduced brain heart infusion agar supplemented with taurocholic acid under anaerobic conditions (37°C) to obtain isolated vegetative colonies. Vegetative cells were cultured or transferred to 70:30 sporulation agar and incubated anaerobically for 4–5 days to induce sporulation. Spores were harvested and stored at 4°C until use.

*Candida auris* (*C. auris*) (BEI Resources NR-52713; strain AKU-2017-385), a multidrug-resistant clinical isolate resistant to amphotericin B and fluconazole, was included as an emerging multidrug-resistant healthcare-associated fungal pathogen, providing a stringent environmental disinfection challenge due to its ability to persist on healthcare surfaces and resist commonly used disinfectants. Single colonies were inoculated into 5 mL of standard fungal growth media (e.g., YPD broth) and incubated at 37°C for 24 h before experimentation.

### Pathogen Quantification

Viral infectivity following UV-C exposure was quantified using a tissue culture infectious dose 50% (TCID50) assay adapted from ASTM E1053 guidelines. After treatment, dried viral inocula were recovered in 10 µL of DMEM supplemented with 10% FBS and incubated for 5 min. Samples were serially diluted 10-fold and inoculated onto Vero E6 cell monolayers seeded in 96- well plates (∼1.5 × 10^4^ cells per well). Following a 1-hour adsorption period at 37°C with 5% CO_2_, the inoculum was removed, fresh DMEM containing 10% heat-inactivated FBS was added, and cultures were incubated for at least 3 days until cytopathic effects (CPE) developed. Viral infectivity was assessed by light microscopy, representative images were acquired using an EVOS M7000 imaging system, and TCID50 values were calculated using the Spearman–Kärber method. Log_10_ reductions were determined relative to matched untreated controls [11].

Viable bacterial, fungal, and spore recoveries following UV-C exposure were quantified by colony-forming unit (CFU) enumeration in accordance with ASTM E1153 and E3135 methodologies. Following treatment, dried inocula were recovered in 10 µL of the appropriate growth medium, serially diluted 10-fold (or 20-fold for spores), and plated onto organism-specific agar media. Plates were incubated under organism-appropriate conditions before colony enumeration. Where technical replicate spot counts were performed, counts were averaged within each independent biological replicate before calculation of log_10_ reductions. Untreated control samples were processed in parallel with UV-C-treated samples and served as the baseline for determination of microbial inactivation. Log_10_ reductions were calculated from the recoverable viable burden of matched untreated and UV-C-treated samples rather than from the nominal inoculum concentration, thereby preventing losses associated with inoculation, drying, surface adherence, and sample recovery from being attributed to UV-C treatment. Percent inhibition was calculated from the corresponding geometric mean recoveries.

### UV-C exposure conditions

Samples were exposed to UV-C under controlled stationary conditions at fixed source-to-surface distances selected to represent practical deployment conditions. Representative high-touch materials were evaluated at 1.5 meters from the UV-C source, whereas stainless-steel surfaces were evaluated at 1.0 meter. Delivered UV-C dose was determined from direct measurements of cumulative UV-C energy (mJ/cm^2^) at the sample location for each exposure condition. At 1.5 meters, measured doses ranged from 2.37 mJ/cm^2^ following a 2-second exposure to 1,300 mJ/cm^2^ following a 7-minute exposure. At 1.0 meter, measured doses ranged from 375 mJ/cm^2^ following a 4-minute exposure to 3,600 mJ/cm^2^ following a 20-minute exposure. Exposure durations were selected according to the anticipated UV-C susceptibility of each organism to encompass the range of reductions required for quantitative analysis. Delivered UV-C dose (mJ/cm^2^) was calculated as: Dose (mJ/cm^2^) = Irradiance (mW/cm^2^) × Exposure Time (seconds). Experimentally measured UV-C doses were used throughout all dose-response analyses, operational dose calculations, and threshold estimations.

### Operational UV-C Dose Analysis

Operational UV-C efficacy was characterized using the experimentally delivered UV-C dose and corresponding observed log_10_ reduction for each pathogen. Where multiple exposure levels provided sufficient dose-response information, interpolation was used to estimate the UV-C dose associated with a defined reduction level within the experimentally observed range. For pathogens evaluated primarily at a single operational dose, the experimentally tested dose and corresponding microbial reduction were reported directly and were not interpreted as minimum effective-dose thresholds.

For adenovirus, the experimentally observed dose-response relationship permitted interpolation of the UV-C dose associated with a 4-log_10_ reduction. The dose-response relationship was constrained to be non-decreasing with increasing UV-C dose using monotonic interpolation, and the dose associated with the 4-log_10_ reduction was estimated within the experimentally evaluated dose range. For the remaining pathogens, experimentally evaluated operational UV-C doses and their corresponding observed microbial reductions were reported directly rather than interpreted as minimum effective doses.

Separately, the multi-dose *C. auris* dataset was used to examine how experimentally determined microbial inactivation could be translated into operational exposure parameters. Dose-level mean log_10_ reductions were constrained to be non-decreasing with increasing UV-C dose using isotonic regression implemented with the pool-adjacent-violators algorithm. A physical anchor of zero UV-C dose and zero log_10_ reduction was included. For this operational modelling analysis, a 2- log_10_ reduction was selected as a defined reference level within the experimentally observed dose-response range. This value was used solely as a reference point for dose-to-time modelling and was not considered a disinfection efficacy threshold, minimum effective dose, or maximum achievable inactivation. The UV-C dose associated with the 2-log_10_ reference level (*D*_2_-log) was estimated by linear interpolation between the adjacent fitted dose levels that bracketed this value.

To translate the estimated *C. auris D*_2_-log into predicted operational exposure times, irradiance at different source-to-surface distances was estimated using the inverse-square relationship. Under this relationship, irradiance (*I*) is proportional to the square of the distance (*r*) from the source: *I_2_ = I_1_(r_1_/r_2_)^2^*, where *I_1_* and *I_2_* are irradiances at source-to-surface distances *r_1_* and *r_2_*, respectively. Using the measured RAY irradiance of approximately 3.0 mJ/cm^2^/second at 1.0 meter, the predicted exposure time required to deliver the estimated dose was calculated as: *t* = (*D*/*I*), where *D* is the target UV-C dose and *I* is the estimated irradiance at the corresponding distance. These distance-dependent exposure times represent theoretical predictions rather than experimentally validated disinfection cycles.

## Statistical Analysis

All experiments were performed using at least three independent biological replicates, with technical replicates included where applicable. Data are presented as mean ± standard deviation (SD) unless otherwise indicated. For the stainless-steel experiments, technical triplicate spot counts were averaged within each independent biological replicate before calculation of log_10_ reductions and statistical analysis. Bacterial, fungal, and spore counts were log_10_-transformed prior to analysis to calculate geometric means and log_10_ reductions in accordance with ASTM recommendations. Viral infectivity was quantified as log_10_ TCID50 values calculated using the Spearman–Kärber method.

All microbial reductions were calculated relative to matched untreated controls. Statistical comparisons were performed only when replicate-level data were available under matched experimental conditions. For experiments in which UV-C-treated samples were compared with matched untreated controls, differences in log_10_-transformed viable recovery were evaluated using two-tailed paired Student’s t-tests, with independent biological experiments serving as the statistical units and technical replicates averaged within each experiment before analysis. This included the viral surface experiments and the stainless-steel experiments for which matched replicate-level controls were available. For comparisons of UV-C efficacy among representative high-touch materials, mean log_10_ reductions were compared using one-way analysis of variance (ANOVA). Where the overall ANOVA was significant, results were interpreted as evidence of differences in the magnitude of the observed reductions among materials rather than differences in the presence or absence of UV-C-mediated inactivation.

Relationships between UV-C dose and microbial inactivation were evaluated by simple linear regression where complete dose-response data were available. Pathogen-specific operational UV-C dose thresholds were estimated by monotonic interpolation of experimentally observed dose-response relationships when sufficient intermediate exposure conditions were available. For organisms evaluated primarily at a single operational UV-C dose (*K. pneumoniae*, *E. faecalis*, and *C. difficile* spores), the experimentally evaluated dose and corresponding microbial reduction were reported directly rather than interpolated. The multi-dose *C. auris* dataset was used for operational modelling because it provided multiple experimentally evaluated UV-C doses spanning the selected 2-log_10_ reference threshold.

Measurements yielding no recoverable organisms were treated as lower-bound estimates corresponding to the assay detection limit. These values were retained for graphical presentation and operational dose determination but were not used to extrapolate dose-response relationships beyond the measurable range. For comparisons among representative materials, results for datasets containing lower-bound estimates should be interpreted as descriptive assessments of the observed reductions.

Dose-response modelling. For *C. auris*, dose-level mean log_10_ reductions were constrained to be non-decreasing with increasing UV-C dose using isotonic regression implemented with the pool-adjacent-violators algorithm. A physical anchor of zero UV-C dose and zero log_10_ reduction was included. For operational modelling, a 2-log_10_ reduction was selected as a defined reference threshold and was not intended to represent a minimum disinfection criterion or maximum achievable inactivation. The UV-C dose associated with this threshold (*D*_2_-log) was estimated by linear interpolation between the adjacent fitted dose levels that bracketed the 2-log_10_ threshold.

Statistical analyses and data processing were performed in Python v3.11 using pandas v2.2.3, NumPy v2.2.6, SciPy v1.15.3, matplotlib v3.10.5, and openpyxl v3.1.5. Statistical significance was defined as *P* < 0.05.

## RESULTS

### Experimental design for UV-C disinfection across representative environmental surfaces

To characterize UV-C dose–response relationships and operational exposure requirements under representative environmental conditions, pathogen viability was assessed following UV-C exposure on a diverse range of high-touch materials, including polystyrene, Kydex, Boltaron, cloth, and synthetic leather, together with healthcare-grade stainless-steel surfaces. Collectively, these materials represent smooth, non-porous and more structurally complex surfaces commonly encountered in healthcare and other high-touch environments.

Disinfection efficacy was quantified as log_10_ reductions relative to matched untreated controls. Untreated control recoveries varied among organisms and materials, reflecting differences in microbial adherence and recovery efficiency, but remained within the quantifiable range required for determination of log_10_ reductions (Supplementary Table S1). Pathogens were evaluated under exposure conditions selected according to the experimental objectives and biological characteristics of each organism. Surface testing was therefore not performed uniformly across all pathogen–material combinations; rather, representative materials were selected to model clinically and environmentally relevant exposure scenarios. Consequently, comparisons between materials are interpreted within the context of individual pathogens rather than as direct cross-surface comparisons. The following sections present pathogen-specific disinfection performance on representative high-touch and healthcare surfaces, followed by an integrated analysis of operational UV-C dose requirements across pathogen classes.

### Dose-dependent inactivation of adenovirus across representative high-touch materials

To evaluate UV-C efficacy against structurally resilient viral pathogens, adenovirus was used as a representative non-enveloped virus and evaluated on polystyrene plastic, Kydex, and Boltaron surfaces. Adenovirus exhibited a clear dose-dependent response, with increasing delivered UV-C dose producing progressively greater reductions in viral infectivity (Figure 2). At a delivered UV-C dose of 78 mJ/cm^2^ (∼60 seconds at 1.5 meters) experimentally tested condition, all evaluated materials achieved reductions of ≥4 log_10_, corresponding to ≥99.99% inhibition (Figure 2A). Lower operational doses (13–26 mJ/cm^2^) produced more modest reductions of approximately 2–3 log_10_ (∼99.0–99.9% inhibition) (Figure 2B). Linear regression demonstrated a significant positive relationship between delivered UV-C dose and log_10_ reduction (equivalent to ∼0.047 log_10_ reduction per second under the measured irradiance conditions; *p* < 0.001), confirming a progressive dose-dependent inactivation response. Mean reductions were comparable across the evaluated materials, and no significant differences in log_10_ reduction were detected among surfaces at the 78 mJ/cm^2^ operational dose (one-way ANOVA, *p* > 0.05). These findings indicate that delivered UV-C dose, rather than surface type, was the primary determinant of adenovirus inactivation under the tested conditions.

**Figure 2.**
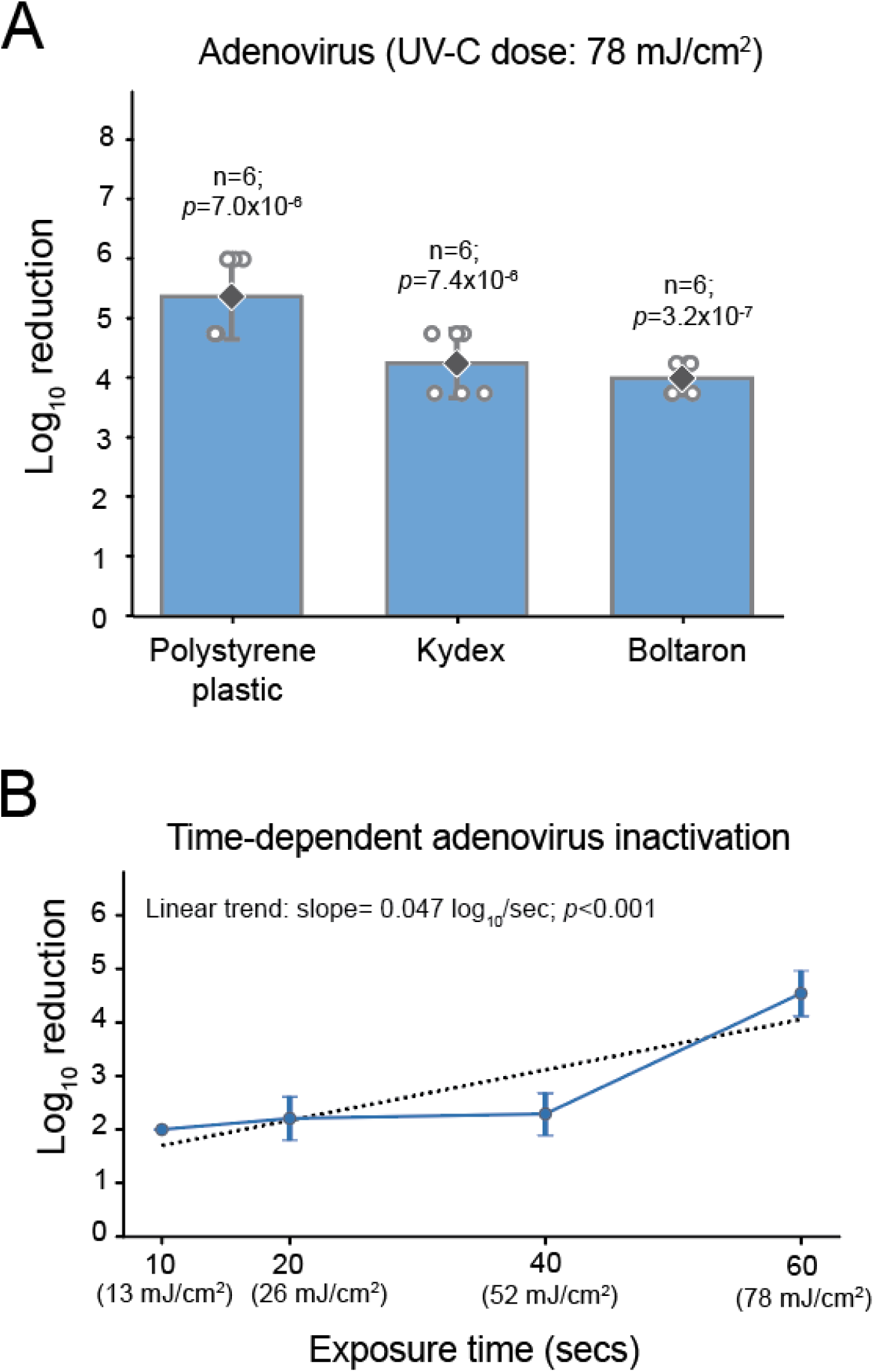
UV-C inactivation of adenovirus across representative high-touch materials. **A**, Log_10_ reduction in adenovirus infectivity following a delivered UV-C dose of 78 mJ/cm^2^ UV-C dose (∼60 seconds at 1.5 meters) from the RAY autonomous UV-C system on representative high-touch materials, including polystyrene plastic, Kydex, and Boltaron. Microbial inactivation is expressed as log_10_ reduction relative to matched untreated controls. Bars represent the mean log_10_ reduction, error bars indicate 95% confidence intervals, open symbols represent independent biological replicates (n = 6 per material), and black diamonds denote the mean. Differences between UV-C-treated samples and their matched untreated controls were evaluated using paired two-tailed Student’s t-tests (*p* values shown), whereas differences in mean log_10_ reduction among materials were evaluated using one-way analysis of variance (ANOVA) (*p* > 0.05). **B**, Dose-dependent adenovirus inactivation following UV-C exposure at 13, 26, 52, and 78 mJ/cm^2^ (corresponding to approximately 10-, 20-, 40-, and 60-second exposures). Points represent the mean log_10_ reduction ± 95% confidence interval pooled across the evaluated materials (n = 3 biological replicates per dose). The dotted line represents the linear regression fitted to replicate-level data, demonstrating a significant positive relationship between delivered UV-C dose and microbial inactivation (*p* < 0.001).

### Rapid inactivation of SARS-CoV-2 across representative materials and variants

To evaluate UV-C efficacy against enveloped viruses, different variants of SARS-CoV-2 were examined across representative high-touch materials. Beta and Gamma variants were evaluated across polystyrene, Kydex, Boltaron, and cloth at a delivered UV-C dose of 13 mJ/cm² (∼10 seconds at 1.5 meters) to assess variant- and material-associated differences in susceptibility (Figure 3A). The original SARS-CoV-2 isolate was evaluated on polystyrene as part of complementary short-exposure experiments designed to characterize UV-C susceptibility at lower doses (Figure 3B). Across the evaluated variant–material combinations, UV-C exposure produced substantial reductions in infectivity, with mean reductions up to approximately 2.0 to 6.1 log_10_, corresponding to approximately 99% to >99.999% inhibition.

**Figure 3.**
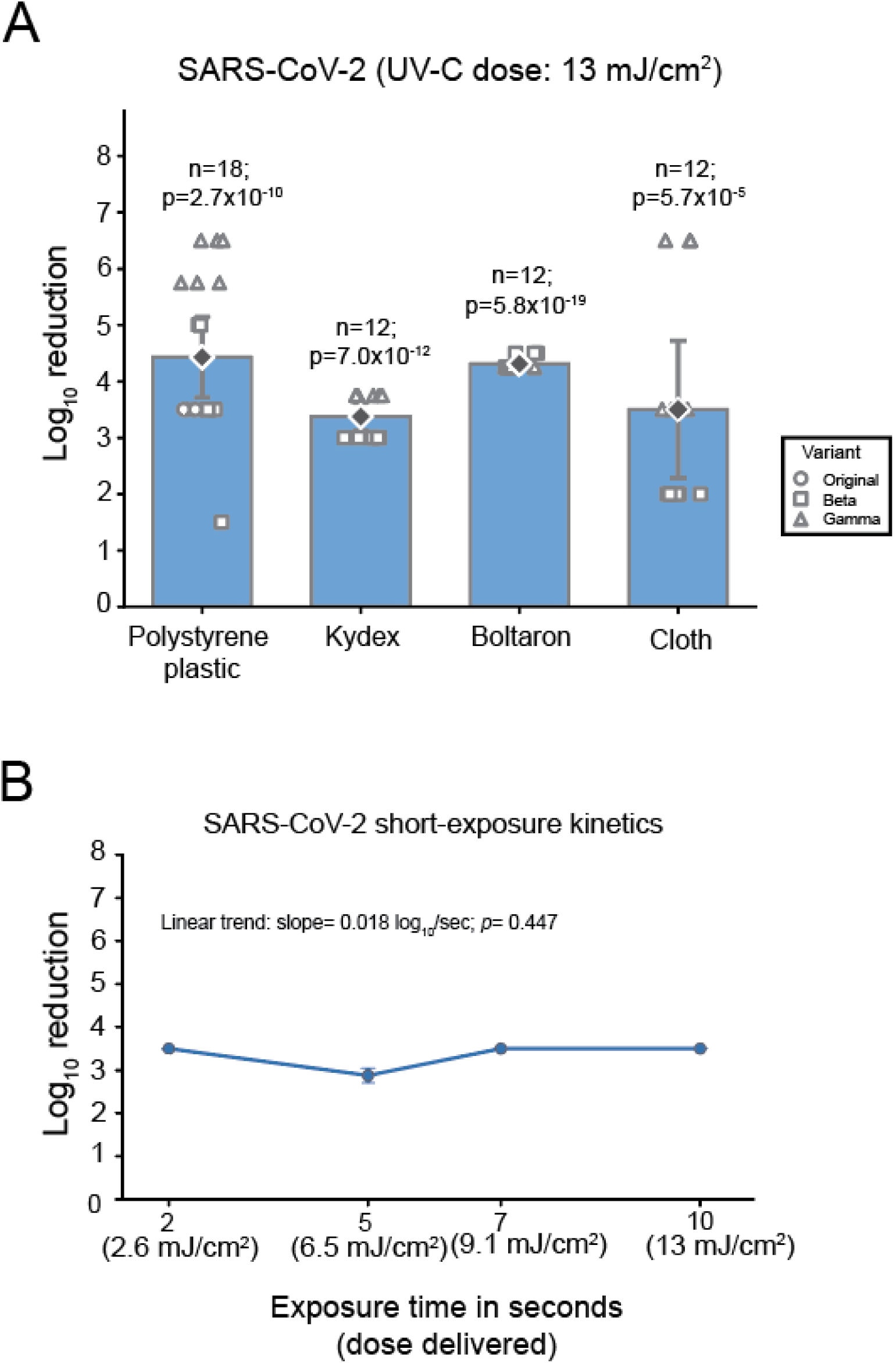
Rapid UV-C inactivation of SARS-CoV-2 across representative high-touch materials and exposure conditions. **A**, Log_10_ reduction in SARS-CoV-2 infectivity following a delivered UV-C dose of 13 mJ/cm² UV-C dose (∼10 seconds at 1.5 meters) from the RAY autonomous UV-C system on representative high-touch materials, including polystyrene plastic, Kydex, Boltaron, and cloth. Microbial inactivation is expressed as log₁₀ reduction relative to matched untreated controls. Bars represent the mean log_10_ reduction, error bars indicate 95% confidence intervals, open symbols represent independent biological replicates, and black diamonds denote the mean. Colored symbols identify the original isolate, Beta, and Gamma variants. Differences between UV-C-treated samples and matched untreated controls were evaluated using paired two-tailed Student’s t-tests (*p* values shown). Differences in inactivation between Beta and Gamma across materials were evaluated using two-way ANOVA to assess the effects of variant, material, and their interaction. Differences among the original isolate, Beta, and Gamma on polystyrene were evaluated using one-way ANOVA followed by Tukey HSD post hoc testing. **B**, Inactivation of the SARS-CoV-2 original isolate on polystyrene plastic following operational UV-C doses of 2.6, 6.5, 9.1, and 13 mJ/cm^2^ (corresponding to approximately 2-, 5-, 7-, and 10-second exposures). Points represent the mean log₁₀ reduction ± 95% confidence interval from six independent biological replicates. The relationship between exposure duration and log₁₀ reduction was evaluated by simple linear regression of replicate- level data.

SARS-CoV-2 variant-specific analysis demonstrated that the magnitude of inactivation differed between Beta and Gamma variants and depended on material. Two-way ANOVA across the four materials identified significant effects of variant (F_1,40_ = 48.76, *p* = 1.98 × 10^-8^), material (F_3,40_ = 10.81, *p* = 2.46 × 10^-5^), and a variant × material interaction (F_3,40_ = 11.19, *p* = 1.82 × 10^-5^). Gamma generally exhibited greater log_10_ reductions than Beta, particularly on polystyrene, Kydex, and cloth, whereas reductions on Boltaron were similar between variants. On polystyrene, where the original isolate, Beta, and Gamma were evaluated under the same 13 mJ/cm^2^ exposure condition, reductions differed significantly among isolates (one-way ANOVA, F_2,15_ = 21.19, *p* = 4.27 × 10^-5^). Tukey post hoc analysis showed significantly greater inactivation of Gamma than Beta (*p* = 0.0002) or the original isolate (*p* = 0.0001), whereas Beta and the original isolate did not differ significantly (*p* = 0.928). These findings indicate that the magnitude of UV-C inactivation varied among SARS-CoV-2 variants, with the extent of these differences influenced by material type.

To further characterize UV-C susceptibility at lower doses, short-exposure experiments were performed on polystyrene using the original isolate (Figure 3B). Rapid viral inactivation (>3.0 log_10_) was evident at the lowest delivered UV-C dose evaluated (2.6 mJ/cm^2^; ∼2 seconds at 1.5 meters), producing a mean reduction of approximately 3.5 log_10_ (99.97% inhibition). Increasing the delivered UV-C dose from 2.6 to 13 mJ/cm^2^ did not produce a significant increase in viral reduction (linear regression, slope = 0.018 log_10_ per second, *p* = 0.447), consistent with rapid attainment of the measurable reduction range under these assay conditions. Accordingly, these data were interpreted as demonstrating substantial SARS-CoV-2 inactivation at very low UV-C doses rather than as defining a minimum effective dose. The variant–material experiments further demonstrated that ≥4-log_10_ reductions were achieved for multiple variant–material combinations at the experimentally evaluated 13 mJ/cm^2^ operational condition.

Collectively, these findings demonstrate that SARS-CoV-2 is highly susceptible to UV-C irradiation, with substantial inactivation occurring at doses as low as 2.6 mJ/cm^2^ and ≥4-log_10_ reductions observed for multiple variant–material combinations at 13 mJ/cm^2^. The magnitude of inactivation varied among variants and materials, with Gamma generally exhibiting greater measured reductions than Beta under the evaluated conditions.

### Surface-dependent bacterial inactivation across representative high-touch materials

To evaluate the influence of representative high-touch materials on bacterial inactivation, the Gram-negative bacterium *P. aeruginosa* and the Gram-positive bacterium *S. aureus* were evaluated on polystyrene plastic, Kydex, Boltaron, and synthetic leather. Following a UV-C dose of 78 mJ/cm^2^ (∼60 seconds at 1.5 meters), *P. aeruginosa* demonstrated highly consistent susceptibility to UV-C irradiation across all tested materials (Figure 4A). Mean reductions ranged from 6.87 to 6.98 log_10_, corresponding to >99.9999% inhibition, with UV-C-treated samples reaching the assay detection limit across all evaluated biological replicates. No significant differences in mean log_10_ reduction were detected among the four material types (one-way ANOVA, F_3,44_ = 0.47, *p* = 0.703). Since all UV-C-treated samples reached the assay detection limit, these comparisons should be interpreted as descriptive assessments of the observed lower-bound reductions rather than precise estimates of material-dependent differences. Nevertheless, comparable reductions were observed on both smooth, non-porous surfaces (polystyrene and Kydex) and more structurally complex materials (Boltaron and synthetic leather), demonstrating consistently high UV-C efficacy across representative high-touch surfaces.

**Figure 4.**
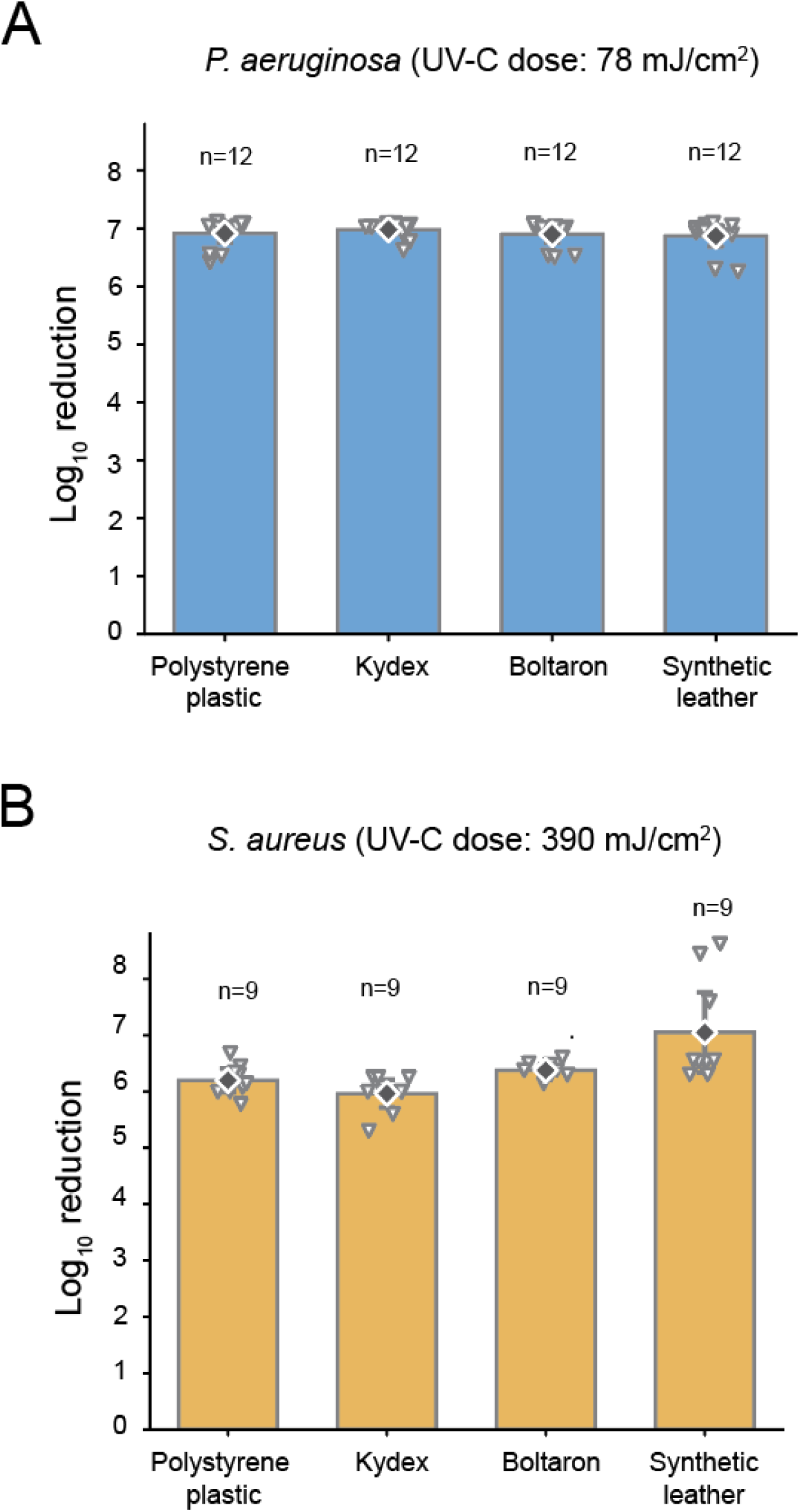
UV-C inactivation of Gram-negative and Gram-positive bacterial pathogens on representative high-touch materials. *P. aeruginosa* (**A**) and *S. aureus* (**B**) were inoculated onto representative high-touch materials, including polystyrene plastic, Kydex, Boltaron, and synthetic leather, and exposed to the indicated delivered UV-C doses under controlled stationary conditions. Microbial inactivation is expressed as log_10_ reduction relative to matched untreated controls. Bars represent the mean log_10_ reduction, error bars indicate 95% confidence intervals, open symbols represent independent biological replicates, and black diamonds denote the mean. For conditions in which no viable bacteria were recovered following UV-C exposure, individual data points represent conservative lower-bound estimates based on the assay detection limit. Differences in mean log_10_ reduction among materials were evaluated using one-way ANOVA (*P. aeruginosa*, F_3,44_ = 0.47, *p* = 0.703; *S. aureus*, F_3,32_ = 7.44, *p* = 0.000651). For *S. aureus*, Tukey HSD post hoc testing identified significantly greater reductions on synthetic leather than on Kydex (adjusted *p* = 0.0005), polystyrene (adjusted *p* = 0.0069), or Boltaron (adjusted *p* = 0.0429); all other pairwise comparisons were not significant.

*S. aureus* also showed substantial susceptibility following a UV-C dose of 390 mJ/cm^2^ (∼5 minutes at 1.5 meters) (Figure 4B). Mean reductions ranged from 5.96 to 7.05 log_10_, equivalent to >99.9998% inhibition across all evaluated materials. Although all surfaces achieved reductions exceeding 5 log_10_, modest differences in mean reduction were observed among materials, with the lowest mean reduction occurring on Kydex and the highest on synthetic leather. Overall differences in mean log₁₀ reduction among materials were statistically significant (one-way ANOVA, F_3,32_ = 7.44, *p* = 0.000651). Tukey HSD post hoc analysis showed significantly greater measured reductions on synthetic leather than on Kydex (adjusted *p* = 0.0005), polystyrene (adjusted *p* = 0.0069), or Boltaron (adjusted *p* = 0.0429), whereas no significant differences were detected among Kydex, polystyrene, and Boltaron (adjusted *p* > 0.05). Despite these differences, all materials achieved >5-log_10_ reductions, indicating that the surface-associated variation reflected differences in the magnitude of already substantial bacterial inactivation rather than reduced UV-C efficacy on any evaluated material.

Collectively, these findings demonstrate substantial bacterial inactivation following the evaluated UV-C exposures across a diverse range of representative high-touch materials. While *S. aureus* exhibited greater variability in measured reductions among materials than *P. aeruginosa*, both organisms showed >5-log₁₀ reductions under the evaluated dose conditions. These findings indicate that substantial bacterial inactivation was maintained across environmentally relevant surfaces differing in composition and structural complexity, although the magnitude of the observed reduction varied by material for *S. aureus*.

### Disinfection of Healthcare-Associated Pathogens on Stainless Steel

To characterize UV-C inactivation of clinically important healthcare-associated pathogens under standardized surface and exposure conditions, stainless-steel coupons were inoculated with four representative pathogens and exposed to defined UV-C doses at a fixed source-to-surface distance of 1.0 meter. Stainless steel was selected because it is widely used on high-touch clinical surfaces, medical equipment, bed rails, carts, and work areas. Delivered UV-C doses were selected to assess organism-specific inactivation while maintaining a consistent surface type and source-to-surface distance across experiments.

*K. pneumoniae* exposed to a UV-C dose of 720 mJ/cm^2^ exhibited the greatest and most consistent reduction in viability (Figure 5A). Across ten independent experiments, the mean reduction was 4.59 log_10_ (95% CI, 3.85–5.34), corresponding to 99.9974% inhibition of recoverable organisms. Nine of ten independent experiments achieved reductions of at least 3 log_10_. Paired comparison of log_10_-transformed viable counts demonstrated a highly significant reduction in bacterial recovery following UV-C exposure (paired two-tailed Student’s t-test, t(9) = 14.02, *p* = 2.02 × 10^-7^).

**Figure 5.**
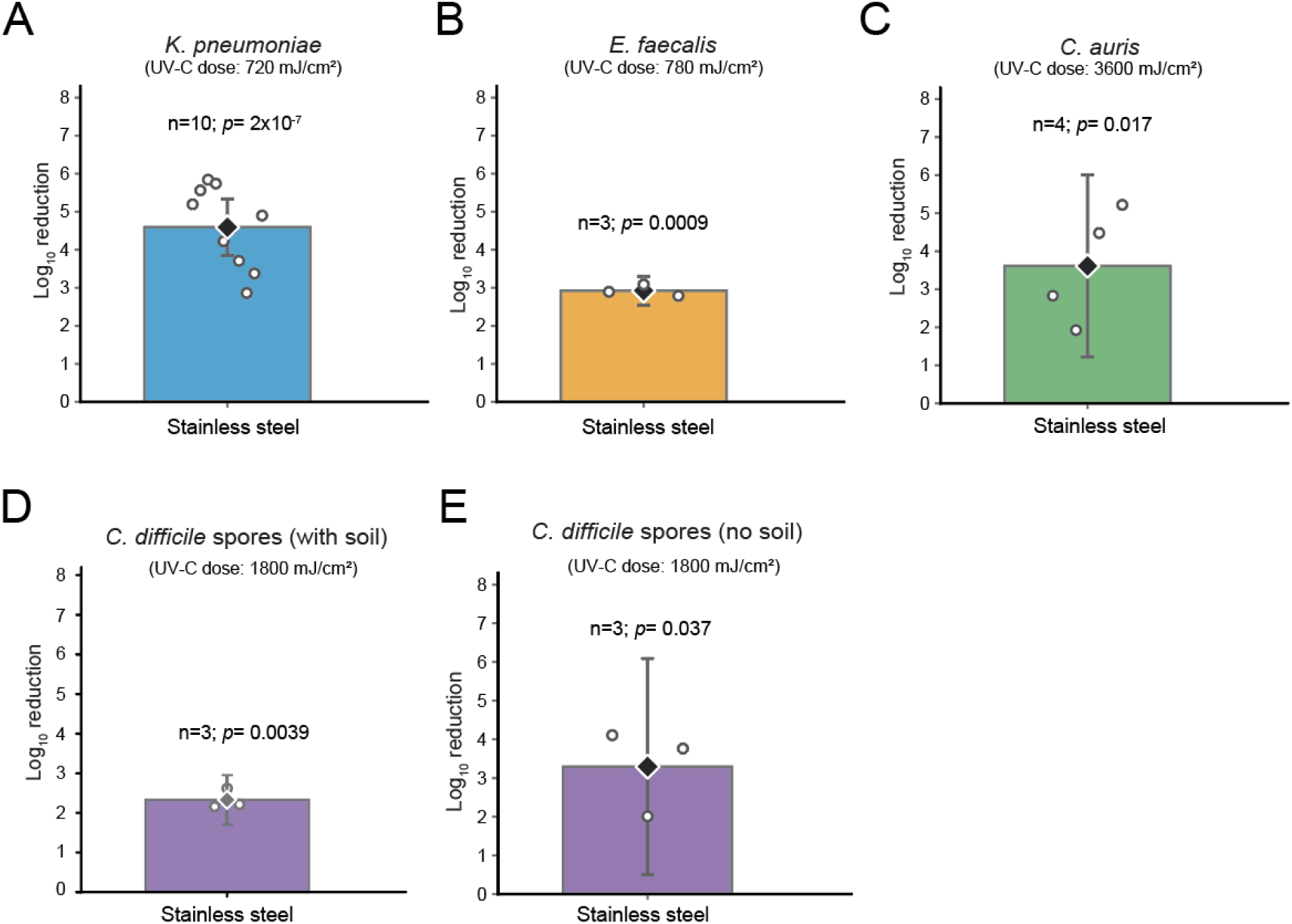
UV-C inactivation of healthcare-associated pathogens on stainless-steel surfaces. Mean log_10_ reduction in viable (**A**) *K. pneumoniae*, (**B**) *E. faecalis*, (**C**) *C. auris*, (**D**) *C. difficile* spores (soil), and (**E**) *C. difficile* spores (no soil) following exposure to the indicated delivered UV-C doses under controlled stationary conditions on stainless-steel coupons at a fixed distance of 1.0 meter. Delivered UV-C doses were 720 mJ/cm^2^ for *K. pneumoniae*, 780 mJ/cm^2^ for *E. faecalis*, and 3600 mJ/cm^2^ for *C. auris*, and 1800 mJ/cm^2^ for *C. difficile* spores. Bars represent the mean log_10_ reduction, error bars indicate the 95% confidence interval, open circles represent independent biological experiments, and filled diamonds denote the calculated group mean. UV- C-treated samples were compared with their matched untreated (no UV-C) controls using two- tailed paired Student’s t-tests, with *p* values shown within each panel. Technical triplicate spot counts were averaged within each independent experiment before calculation of log_10_ reductions.

*E. faecalis* exposed to 780 mJ/cm^2^ demonstrated a mean reduction of 2.92 log_10_ (95% CI, 2.55– 3.30) across three independent experiments, equivalent to 99.8805% inhibition (Figure 5B). UV-C exposure significantly reduced viable recovery relative to matched untreated controls (paired two-tailed Student’s t-test, t(2) = 33.34, *p* = 0.000898).

For *C. auris*, exposure to 3600 mJ/cm^2^ produced a mean reduction of 3.61 log_10_ (95% CI, 1.22– 6.01) across four independent experiments, corresponding to 99.9756% inhibition (Figure 5C). UV-C exposure significantly reduced viable fungal recovery relative to matched untreated controls (paired two-tailed Student’s t-test, t(3) = 4.80, *p* = 0.0172).

*C. difficile* spores exposed to 1800 mJ/cm^2^ exhibited a mean reduction of 2.33 log_10_ (95% CI, 1.71–2.95) across three independent experiments, corresponding to 99.53% inhibition (Figure 5D). UV-C exposure significantly reduced recoverable spore counts relative to matched untreated controls (paired two-tailed Student’s t-test, t(2) = 16.06, *p* = 0.0039). In additional experiments performed without the 10% FBS soil challenge, the same UV-C exposure produced a numerically greater mean reduction of 3.29 log_10_ (95% CI, 1.71–6.09), corresponding to 99.9489% inhibition (Figure 5E). UV-C exposure significantly reduced recoverable spore counts relative to matched untreated controls (paired two-tailed Student’s t-test, t(2) = 5.07, *p* = 0.0368), although the wide confidence interval indicated appreciable between-experiment variability. The increased efficacy in the absence of a soil challenge is consistent with previous evidence that organic material can attenuate UV-C-mediated inactivation of *C. difficile* spores. Since the study was not designed as a formal comparison of soil conditions, this difference was interpreted descriptively.

Overall, UV-C exposure significantly reduced viable recovery of all four healthcare-associated pathogens tested on stainless steel. Mean inactivation ranged from 2.33 to 4.59 log_10_, corresponding to 99.53% to 99.9974% inhibition, with particularly strong and reproducible activity against *K. pneumoniae*. Substantial reductions were also observed for *E. faecalis*, *C. auris*, and the comparatively resistant spores of *C. difficile*. These findings demonstrate substantial microbial inactivation on stainless-steel surfaces across the defined UV-C dose conditions evaluated.

### Integrated Analysis of Pathogen-Specific UV-C Dose and Exposure-Time Requirements

To compare UV-C exposure requirements across pathogen classes, experimentally evaluated conditions were integrated according to delivered UV-C dose and corresponding exposure time (Figure 6). The evaluated UV-C exposures varied substantially among the pathogens evaluated, spanning from a few mJ/cm^2^ delivered within seconds for SARS-CoV-2 to 3600 mJ/cm^2^ delivered over 20 minutes for *C. auris*. Since pathogens were evaluated under different experimental conditions, including different surfaces and source-to-surface distances, these values represent experimentally evaluated UV-C exposure conditions under the tested geometries rather than direct measures of intrinsic UV-C susceptibility.

**Figure 6.**
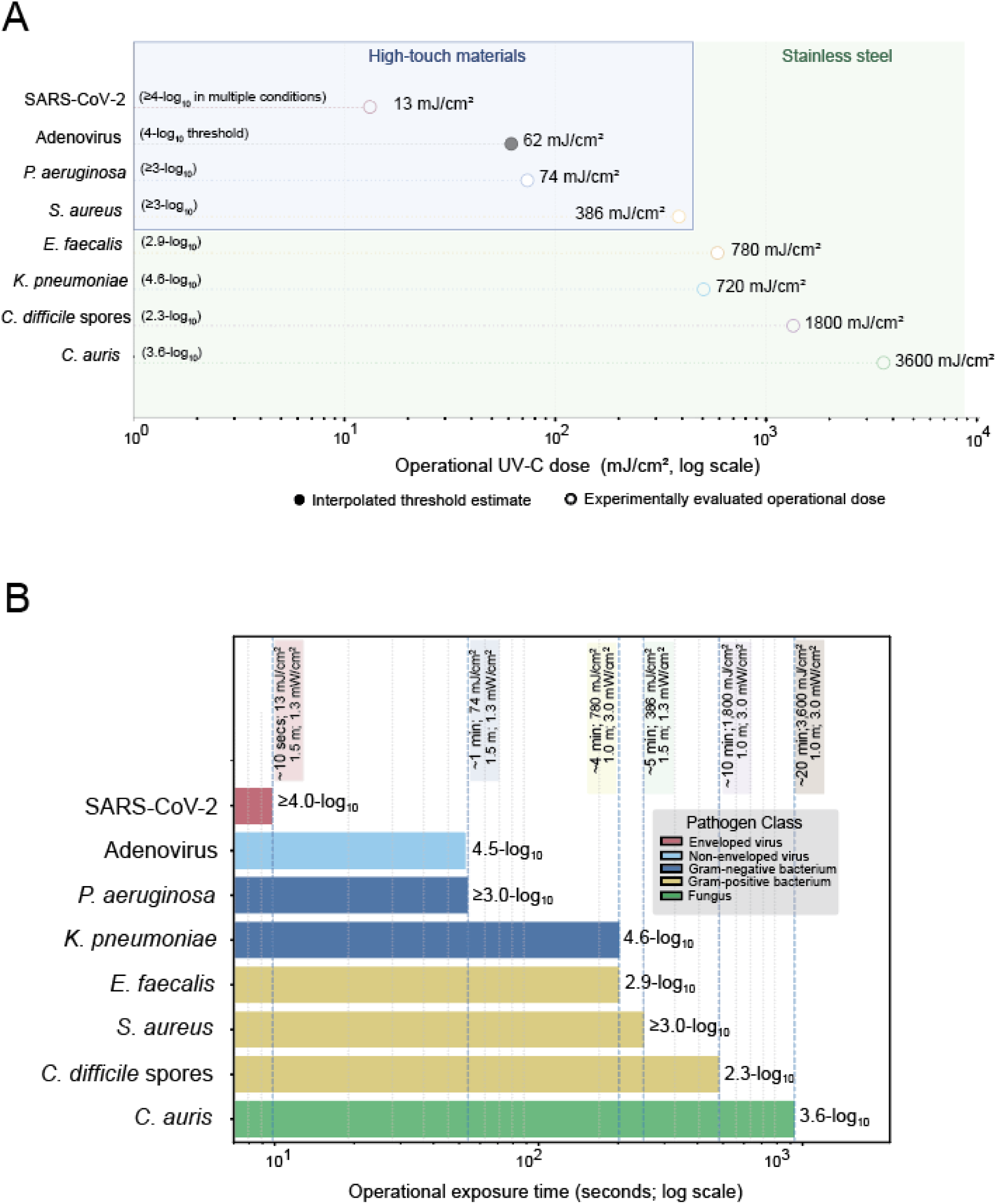
Operational UV-C dose and exposure time requirements for clinically relevant pathogens. A, Experimentally evaluated UV-C doses associated with microbial inactivation across the pathogen classes and surface conditions examined. High-touch materials were evaluated at approximately 1.5 meters from the UV-C source, whereas healthcare-relevant stainless-steel coupons were evaluated at approximately 1.0 meter. For adenovirus, the 4-log_10_ operational dose threshold was estimated by monotonic interpolation of the experimentally observed dose-response relationship. For all other pathogens, symbols represent experimentally evaluated operational UV-C doses and their corresponding observed microbial reductions rather than estimates of minimum effective dose. **B**, Corresponding exposure times associated with the UV-C doses shown in panel A. Labels indicate the microbial reduction observed at each operational exposure; ≥ denotes conditions for which the displayed reduction criterion was achieved or exceeded.

SARS-CoV-2 exhibited substantial inactivation at the lowest UV-C doses evaluated, with >3- log_10_ reduction observed at 2.6 mJ/cm^2^ (∼2 seconds at 1.5 meters) and ≥4-log_10_ reductions observed for multiple variant–material combinations at 13 mJ/cm^2^ (∼10 seconds). Adenovirus exhibited a progressive dose-response relationship, permitting monotonic interpolation of the dose associated with a 4-log_10_ reduction, estimated at approximately 62 mJ/cm^2^ (∼48 seconds at 1.5 meters). The first experimentally evaluated adenovirus condition exceeding this reduction was 78 mJ/cm^2^ (∼60 seconds).

Vegetative bacteria were evaluated at intermediate operational exposures. *P. aeruginosa* and *S. aureus* demonstrated substantial inactivation following experimentally delivered doses of 78 and 390 mJ/cm^2^, respectively, on representative high-touch materials at 1.5 meters. On stainless steel at 1.0 meter, 720 mJ/cm^2^ produced a mean 4.59-log_10_ reduction in *K. pneumoniae*, whereas 780 mJ/cm^2^ produced a mean 2.92-log_10_ reduction in *E. faecalis*. The highest operational doses were evaluated for *C. difficile* spores and *C. auris*, for which 1800 and 3600 mJ/cm^2^ produced mean reductions of 2.33 and 3.61 log_10_, respectively.

Collectively, the experimentally evaluated conditions demonstrate a broad range of UV-C exposures associated with microbial inactivation across the pathogen classes studied (Figure 6). With the exception of the interpolated adenovirus value, these doses should be interpreted as experimentally evaluated operational conditions and their corresponding observed reductions rather than estimates of minimum effective dose.

### Translation of UV-C dose to predicted operational exposure times

To illustrate how experimentally determined UV-C efficacy could be translated into operational exposure times, a 2-log_10_ reduction was selected as a standardized reference threshold within the experimentally observed dose-response range. This threshold was used as a defined reference point for dose-to-time calculations and was not intended to represent a minimum disinfection criterion or the maximum achievable inactivation. Modelling was restricted to *C. auris* because its multi-dose dataset bracketed this threshold, permitting estimation within the experimentally evaluated dose range rather than by extrapolation. An estimated UV-C dose of approximately 1.41 J/cm^2^ was associated with this threshold (Figure 7A). This dose was then used to estimate exposure times at varying source-to-surface distances. Since UV-C irradiance decreases approximately with the square of distance, increasing the distance from the source reduces irradiance and correspondingly increases the time required to accumulate the same UV-C dose. For example, doubling the distance theoretically reduces irradiance 4-fold and therefore requires approximately four times longer to deliver the same UV-C dose. Conversely, decreasing the distance from 1.0 meter to 0.5 meter theoretically increases irradiance 4-fold, reducing the predicted exposure time from approximately 7.8 to 2.0 minutes. Using an irradiance of approximately 3.0 mJ/cm^2^/second at 1.0 meter and the inverse-square relationship *I_2_ = I_1_(r_1_/r_2_)^2^*, the predicted time required to deliver 1.41 J/cm^2^ was approximately 2.0 minutes at 0.5 meter, 4.4 minutes at 0.75 meter, 7.8 minutes at 1.0 meter, 12.3 minutes at 1.25 meters, 17.6 minutes at 1.5 meters, 24.0 minutes at 1.75 meters, and 31.4 minutes at 2.0 meters (Figure 7B). Thus, decreasing source-to-surface distance substantially reduced the predicted exposure time required to deliver a UV-C dose associated with a 2-log_10_ reduction.

**Figure 7.**
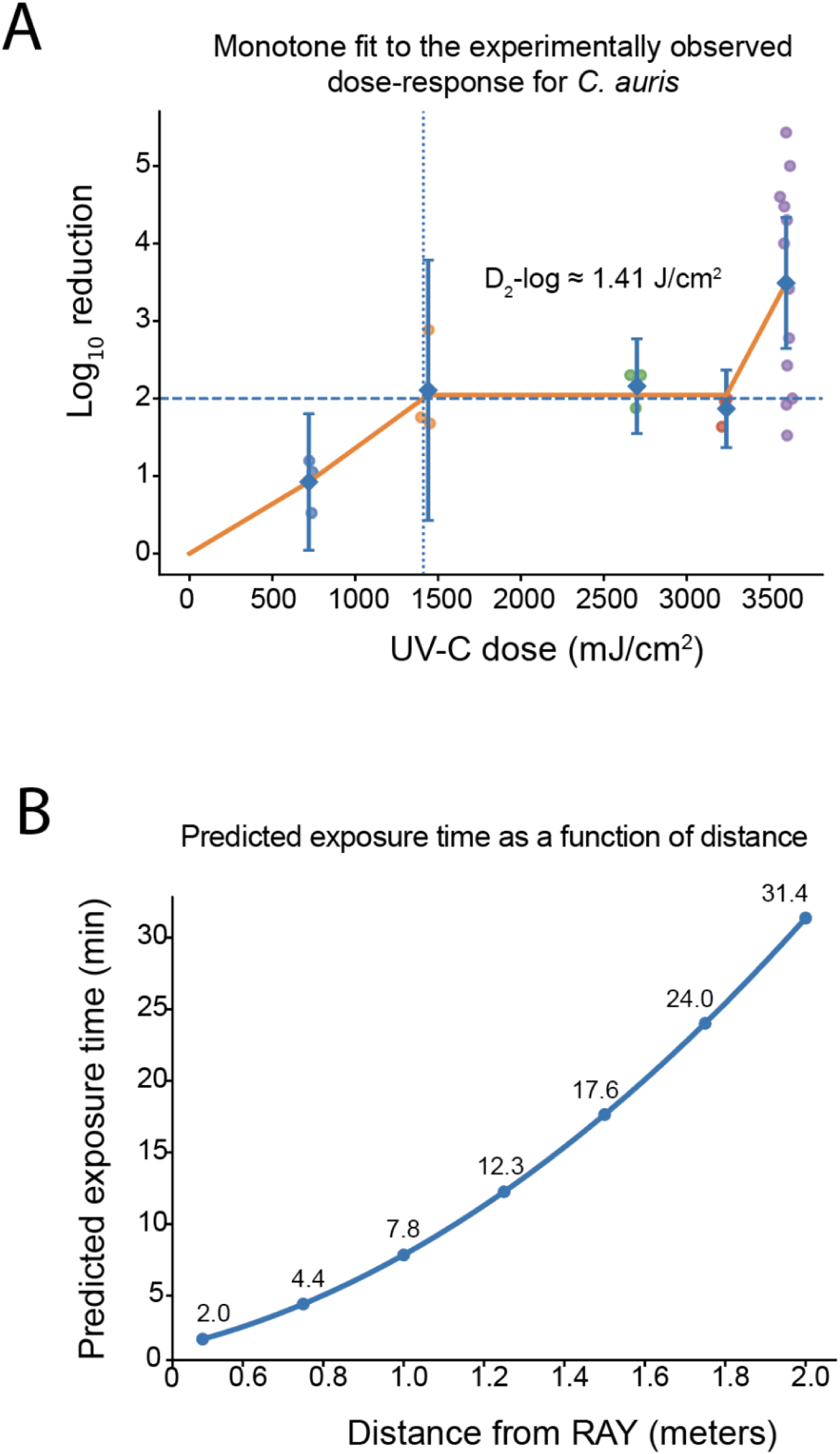
UV-C dose–response relationship for C. auris and predicted exposure times as a function of source-to-surface distance. **A**, Log_10_ reduction in *C. auris* on stainless steel following exposure to increasing UV-C doses. Individual biological replicates and mean values with 95% confidence intervals are shown. For operational modelling, a 2-log_10_ reduction was selected as a standardized reference threshold within the experimentally observed dose-response range and was not intended to represent the maximum achievable inactivation. The estimated UV-C dose associated with this threshold (*D*_2_-log) was approximately 1.41 J/cm^2^. **B**, Predicted exposure time required to deliver the estimated *D*_2_-log at increasing source-to-surface distances. Times were calculated from an irradiance of approximately 3.0 mJ/cm^2^/second at 1.0 meter using the inverse-square relationship, *I_2_ = I_1_(r_1_/r_2_)^2^*, and *t* = *D*/*I*. Predicted exposure time increased with the square of distance, from approximately 2.0 minutes at 0.5 meter to 31.4 minutes at 2.0 meters. Distance-based values represent theoretical dose-to-time estimates rather than experimentally validated disinfection cycles.

## DISCUSSION

In this study, experimentally quantified UV-C doses produced substantial microbial inactivation across diverse pathogen classes and representative high-touch and healthcare surfaces. UV-C inactivation is mediated primarily by photochemical damage to microbial nucleic acids, including the formation of pyrimidine dimers and related lesions that impair replication and infectivity[9]. The broad activity observed across enveloped and non-enveloped viruses, Gram-negative and Gram-positive bacteria, a fungal pathogen, and bacterial spores is therefore consistent with the fundamental mechanism of germicidal UV-C irradiation.

The material panel was selected to represent operational exposure scenarios rather than to provide an exhaustive comparison of every pathogen on every surface. Polystyrene, Kydex, Boltaron, cloth, and synthetic leather captured a range of smooth, composite, and porous high-touch materials, whereas stainless steel represented a standardized healthcare surface with a well-established role in environmental transmission [1–3]. This design extends UV-C evaluation beyond idealized laboratory substrates and addresses material properties that can influence delivered dose, including surface topology, porosity, reflectivity, and local shielding [1,2,4,6,7]. Accordingly, the findings provide an operationally relevant assessment of UV-C performance while recognizing that comparisons between organisms tested on different materials or at different distances do not constitute direct measurements of intrinsic UV-C susceptibility.

The viral data demonstrate the marked difference in operational exposure requirements between enveloped and non-enveloped viruses. SARS-CoV-2 was inactivated within seconds across the evaluated materials, consistent with previous reports of efficient UV-C-mediated inactivation of surface-dried SARS-CoV-2 [6,12]. Adenovirus required greater cumulative exposure to achieve comparable reductions, in agreement with its greater environmental stability and generally higher tolerance to UV irradiation [13–15]. The lack of a significant increase in SARS-CoV-2 inactivation across the evaluated UV-C dose range (∼2.6–13 mJ/cm^2^) likely reflects rapid attainment of the assay’s measurable reduction range rather than an absence of additional photochemical damage at higher doses.

The bacterial experiments further demonstrated that effective operational conditions depend on both organism and surface context. *P. aeruginosa* showed consistently high inactivation across all evaluated high-touch materials following delivery of approximately 78 mJ/cm^2^, with little evidence of a material-dependent effect. *S. aureus* also showed substantial reductions across all materials following delivery of approximately 390 mJ/cm^2^, although the magnitude of inactivation varied significantly by surface. This contrast is compatible with the influence of organism-specific physiology and surface-mediated shielding, but the present design does not isolate the relative contribution of each factor [15,16]. The results therefore support robust bacterial inactivation under the tested conditions while emphasizing the value of validating operational doses on representative materials rather than extrapolating from a single standardized substrate.

The stainless-steel experiments extend the study to clinically important healthcare-associated pathogens under standardized operational exposure conditions. Significant reductions in viable recovery were observed for *K. pneumoniae*, *E. faecalis*, *C. auris*, and *C. difficile* spores relative to matched untreated controls. The particularly consistent response of *K. pneumoniae* demonstrates strong activity against a carbapenem-resistant *Enterobacterales* strain, whereas the *E. faecalis* findings show substantial inactivation of a vancomycin-resistant isolate. Together, these results demonstrate that defined UV-C exposures can substantially reduce viable multidrug-resistant bacterial pathogens on stainless steel under standardized experimental conditions.

The *C. auris* findings are notable because this emerging multidrug-resistant fungal pathogen can persist on healthcare surfaces and patient-care equipment, creating challenges for environmental decontamination [17–19]. Previous studies have demonstrated that UV-C can substantially reduce viable *C. auris*, but efficacy is strongly influenced by exposure time, source-to-surface distance, strain, exposure geometry, and organic soil [20–22]. At the highest UV-C dose evaluated in the present study (3.6 J/cm^2^), a mean 3.61-log_10_ reduction was observed despite the standardized 10% FBS organic soil challenge. This magnitude of inactivation is consistent with previous UV-C studies demonstrating substantial reductions in *C. auris* under appropriately delivered exposure conditions. Notably, Rutala et al. reported 4.57-log_10_ reduction of *C. auris* in the presence of a fetal calf serum organic challenge under direct line-of-sight UV-C exposure, compared with 2.41-log_10_ under indirect exposure, demonstrating the importance of exposure geometry [22]. More recently, Vincent et al. observed a 2.6-log_10_ reduction under a low-soil condition but only a 1.0-log_10_ reduction under a 5% soil condition during far-UV-C exposure in hospital patient rooms, further demonstrating that organic material can substantially influence measured *C. auris* inactivation [20]. Collectively, these findings support the substantial UV-C-mediated inactivation observed here while emphasizing that comparisons among UV-C studies must account for differences in delivered dose, wavelength, organic load, source-to-surface distance, and exposure geometry.

Importantly, the multi-dose *C. auris* dataset also provided an opportunity to examine how experimentally determined microbial inactivation could be translated into practical operational parameters. Rather than attempting to derive dose-response thresholds for all organisms, for which the number and distribution of evaluated doses differed, modelling was restricted to *C. auris* because multiple experimentally evaluated doses spanned a defined 2-log_10_ reduction threshold. A 2-log_10_ reduction was selected as a reference point for operational modelling and was not intended to represent either a minimum disinfection criterion or the maximum inactivation achievable under the evaluated UV-C exposure conditions. Interpolation within the experimentally observed *C. auris* response estimated that approximately 1.41 J/cm^2^ was associated with this reference reduction. The importance of this analysis is therefore not the selection of a 2-log_10_ endpoint itself, but the demonstration that experimentally determined microbiological efficacy can be converted into a quantitative UV-C dose target and subsequently into predicted operational exposure times.

This translation also illustrates the strong influence of source-to-surface distance on UV-C delivery. Using the measured irradiance of approximately 3.0 mJ/cm^2^/second at 1.0 meter and an inverse-square relationship, the estimated 1.41 J/cm^2^ *C. auris* reference dose corresponded to predicted exposure times of approximately 2.0 minutes at 0.5 meter, 7.8 minutes at 1.0 meter, 17.6 minutes at 1.5 meters, and 31.4 minutes at 2.0 meters. Under an ideal inverse-square relationship, irradiance decreases in proportion to the square of distance; consequently, doubling the source-to-surface distance decreases irradiance approximately 4-fold and requires approximately four times longer to accumulate the same UV-C dose. Conversely, reducing the distance from 1.0 to 0.5 meter theoretically increases irradiance 4-fold and reduces the predicted exposure time from approximately 7.8 to 2.0 minutes. This relationship has direct implications for UV-C dose delivery because source positioning can substantially alter the time required to accumulate a defined microbiologically relevant dose. Reducing source-to-surface distance can therefore shorten the exposure duration required to achieve a specified cumulative UV-C dose, providing a quantitative basis for optimizing source positioning during automated disinfection.

These distance-based estimates should not, however, be interpreted as experimentally validated disinfection cycles. The inverse-square relationship provides an idealized physical model and assumes that irradiance varies principally as a function of source-to-surface distance. Because the RAY platform incorporates an extended UV-C source rather than an ideal point source, and because UV-C delivery in real environments is additionally influenced by surface orientation, line-of-sight, object shadowing, reflectance, room geometry, and lamp configuration, the inverse-square model represents an idealized approximation. Thus, the model is most appropriately viewed as a demonstration of how an experimentally derived microbiological dose target can be translated into predicted operational parameters. Direct irradiance measurement or dosimetry at target surfaces remains preferable when defining exposure conditions in complex environments.

The inclusion of *C. difficile* spores broadens the relevance of the study to one of the most difficult environmental targets in healthcare. Bacterial spores are protected by a dehydrated core, multilayered coat structures, and DNA-associated protective proteins that collectively increase tolerance to physical and chemical stress. Despite these protective adaptations, delivery of 1800 mJ/cm^2^ under the standardized 10% FBS organic soil challenge produced a mean 2.33-log₁₀ reduction (99.53%) in recoverable spores on stainless steel. Although variability in UV-C susceptibility can arise from heterogeneity among *C. difficile* spores, including strain- and spore- dependent differences in exosporium structure, surface hydrophobicity, and adherence to stainless steel, these factors were not directly evaluated in the present study. Nevertheless, the significant reduction relative to matched untreated controls demonstrates substantial UV-C-mediated inactivation of *C. difficile* spores under an organic soil challenge and supports further evaluation of dose-controlled UV-C as an adjunct to manual cleaning and chemical disinfection for reducing environmental contamination by spore-forming pathogens.

A central contribution of this study is the quantitative integration of delivered UV-C dose, microbial inactivation, surface context, and operational exposure conditions across diverse pathogen classes. For most pathogens, the experimentally evaluated doses and corresponding reductions establish effective exposure conditions under the tested geometry but should not be interpreted as minimum effective doses or as direct measures of intrinsic UV-C susceptibility across organisms. The *C. auris* dataset provides a complementary proof-of-concept demonstrating how a sufficiently resolved dose-response series can be used to define a microbiological reference dose and translate that dose into predicted exposure times. Together, these approaches distinguish experimentally demonstrated efficacy from operational modelling while providing a framework through which additional pathogen-specific dose-response datasets could subsequently be incorporated.

These findings are directly relevant to autonomous UV-C deployment. The stationary experiments at defined distances were designed to provide reproducible quantification of delivered dose while minimizing variability introduced by movement, room geometry, and dynamic positioning. The experimentally evaluated dose conditions established here could therefore serve as microbiologically grounded targets for autonomous systems that adjust source position and dwell time according to cumulative UV-C exposure. The *C. auris* modelling further illustrates the potential value of proximity optimization, as reducing source-to-surface distance can substantially shorten the predicted time required to deliver a defined UV-C dose. Translation to real-world autonomous operation will nevertheless depend on line-of-sight, surface orientation, obstruction, room geometry, lamp output, environmental conditions, and direct verification of dose delivery at target surfaces [23].

Several limitations should be considered. First, experiments were performed under controlled stationary exposure conditions rather than in occupied or simulated healthcare environments. This design was intentional, allowing precise quantification of delivered UV-C dose and direct assessment of pathogen responses to defined operational exposures while minimizing variability arising from room geometry, shadowing, surface orientation, and robot movement. Their translation to room-scale deployment will depend on factors including room geometry, object shadowing, surface orientation, reflectance, and dynamic positioning. Similarly, the distance-dependent exposure times modelled for *C. auris* were derived using an idealized inverse-square relationship and should be considered theoretical predictions rather than validated clinical disinfection cycles. Further studies incorporating direct surface dosimetry during autonomous operation in representative healthcare environments will be important for determining how closely these predictions correspond to delivered doses under real-world conditions. The controlled efficacy, dose-response, and operational exposure data established here nevertheless provide a quantitative benchmark for configuring and evaluating autonomous UV-C deployment under such conditions.

In summary, experimentally quantified UV-C exposures produced substantial microbial inactivation across viruses, vegetative bacteria, *C. auris*, and *C. difficile* spores on representative high-touch and healthcare surfaces. The study establishes experimentally evaluated UV-C exposure conditions across diverse pathogen classes and materials and, using *C. auris* as a proof-of-concept, demonstrates how a resolved dose–response relationship can be translated into distance-dependent exposure-time predictions. The marked effect of source-to-surface distance on predicted treatment time further highlights the potential advantage of autonomous positioning and dose-aware route optimization. Together, these findings provide a quantitative framework linking delivered UV-C dose, microbial reduction, and operational deployment while defining the experimental and environmental factors that must be considered when translating controlled efficacy data into real-world infection-prevention workflows.

## Author Contributions

Study conceptualization, S.D.B.; methodology, S.D.B.; formal analysis, I.W., N.V., L-V.S-V., E.W., P.S-R., N.J.F., K.A.F., and S.D.B.; investigation, I.W., N.V., L-V.S-V., E.W., P.S-R., N.J.F., K.A.F., and S.D.B.; resources, I.W., N.V., L-V.S-V., E.W., K.R., P.S-R., E.R.I., N.J.F., K.A.F., and S.D.B.; data curation, I.W., N.V., L-V.S-V., E.W., P.S-R., N.J.F., K.A.F., and S.D.B.; writing—original draft preparation, S.D.B.; writing—review and editing, I.W., N.V., L-V.S-V., E.W., P.S-R. and S.D.B.; supervision, S.D.B.; project administration, S.D.B.; funding acquisition, S.D.B. All authors have read and agreed to the published version of the manuscript.

